# Cryptsim: Modeling the evolutionary dynamics of the progression of Barrett’s esophagus to esophageal adenocarcinoma

**DOI:** 10.1101/323485

**Authors:** Diego Mallo, Rumen Kostadinov, Luis Cisneros, Mary K. Kuhner, Carlo C. Maley

## Abstract

To alleviate the over-diagnosis and overtreatment of premalignant conditions we need to predict their progression to cancer, and therefore, the dynamics of an evolutionary process. However, monitoring evolutionary processes *in vivo* is extremely challenging. Computer simulations constitute an attractive alternative, allowing us to study these dynamics based on a set of evolutionary parameters.

We introduce *CryptSim*, a simulator of crypt evolution inspired by Barrett’s esophagus. We detail the most relevant computational strategies it implements, and perform a simulation study showing that the interaction between neighboring crypts may play a crucial role in carcinogenesis.

## I. Introduction to the type of problem in cancer

Pre-malignant conditions are changes in the normal tissue structure-usually characterized by dysplasia or benign neoplasia-that increase the risk of developing a specific kind of cancer. These conditions are generally detected via screening, and their association with cancer fosters their surveillance and aggressive treatment. However, their probability of progression to invasive cancer is often low (e.g., 0.12% of patients with Barrett’s esophagus (BE) without dysplasia progress to esophageal adenocarcinoma every year [1]), and therefore most actions carried out on patients with these conditions constitute over-treatment. To reduce over-treatment we need to improve our risk stratification procedures, which means we must improve our ability to predict the progression of pre-malignant conditions to invasive cancers. Cancer is a disease of the somatic genome in which somatic cells mutate and evolve. Therefore, predicting cancer progression implies predicting evolution. While rough estimates of somatic evolutionary parameters are available (see [3] for a summary), these are not enough to fully understand or predict cancer progression. Studying evolutionary processes *in vivo* is extremely difficult due to economic, ethical, temporal and sampling-related limitations. Moreover, because somatic evolution is a stochastic process, we may require impractically large sample sizes to detect regularities. Computer simulations are an attractive alternative and complementary approach to surveillance of patient cohorts. Simulations allow us to perform as many replicates as are needed and eliminate sampling limitations. There is, however, a combinatorial explosion in the number and combinations of parameters, and so we restrict the investigation of the model to a set of *a priori* most relevant parameters.

Barrett’s esophagus (BE) is a pre-malignant condition that increases the risk of progression to esophageal adenocarcinoma [4]. The BE epithelium is divided into crypts, well-like structures with a few stem cells near the bottom. Crypts are relatively homogeneous units of selection that accumulate mutations, divide and die. High genetic diversity among crypts at initial endoscopy is associated with progression [5,6], and clonal expansions via crypt divisions have been observed [7,8]. However, little is known about the dynamics of this process or the relative importance and role of the different evolutionary parameters that drive progression.

In this highlight, we introduce *CryptSim* [9]-a simulator of crypt evolution inspired by BE. Using it, we show that the interaction between neighboring crypts plays a key role in carcinogenesis of neoplasms that occur in crypt-based two-dimensional tissues, constituting a promising target for cancer prevention.

## II. Illustrative Results of application of Methods

We performed a series of simulation studies to assess the effect of different evolutionary and crypt-biology parameters in carcinogenesis (see Table 1 for a list of simulation parameters). It is unclear if a dividing crypt can replace a neighbor or if it must wait for a neighbor to die before it can divide. We model this with a parameter for the probability of replacing a neighbor crypt. The value of this parameter dramatically affected progression to cancer. Reducing the probability that a crypt could replace a neighbor reduced the probability of progression, and zero probability of replacing a neighbor completely stopped progression (Fig. 1).

The implications of this finding are twofold. On the one hand, it shows the need for a better understanding of basic crypt biology to better model and predict cancer progression. Specifically, little is known about the process and dynamics of tissue homeostasis at the crypt level (e.g., control of replication, replication rate, and death rate). On the other hand, it highlights the key role of spatial competition in cancer progression. Progression to cancer requires the acquisition of a few advantageous mutations (i.e., driver mutations) in (at least) one clone in order to acquire the hallmarks of cancer [10]. Large population sizes increase the rate of fitness increases by mutations and natural selection. However, in most tissues population sizes are limited by spatial competition.

**TABLE I.** SIMULATION PARAMETERS

| Parameter | Value |
| --- | --- |
| N. selective loci, $m$ | 5 reproductive selection loci |
| Mutation rate, $\mu$ | $10^{-5}$ mutations/locus/replication |
| Selective coefficients, $r$ | 0.3, 0.66, 1.13, 1.79, 2.92 |
| Mutation model, $a$ | Time-dependent |
| Crypt division time, $b$ | 50 days |
| Tissue size, $h \times w$ | 300 x 300 crypts |

The replacement of neighboring crypts included in our model facilitates somatic evolution and constitutes a promising target for cancer prevention. Although high tissue renewal rates (i.e., high replication and death rates) and reductions in crypt size are additional mechanisms that could provide additional space for evolution to act, they would require unrealistic parameter values to allow cancer progression.

## III. Quick Guide to the Methods

*Cryptsim’s* model is composed of three main constituents: an agent-based model, a two-dimensional spatial model, and a clone tree. The agent-based model describes the evolution of crypts through time. Using crypts rather than cells as our basic unit allows us to keep computational requirements of realistic tissue sizes within reason and is based on the fact that the small number of stem cells in a crypt are thought to be rapidly homogenized by genetic drift. Each crypt *i* has associated a genome *X*_*i*_ composed of *n* neutral and *m* selective loci *l*=*m*+*n*. Crypts can undergo four different events, each controlled by a rate: replication (*b*), death (*d*), mutation of neutral loci (*v*), and mutation of selective loci (μ). All events can act freely except replication, which requires free space in the neighborhood of the dividing crypt. If there is none, the new crypt can replace a neighbor at random with probability *p*. While most simulation parameters are constant and shared for all crypts for each simulation run, *X*_*i*_, *b*_*i*_, and *d*_*i*_ are private to the crypts and change with the accumulation of mutations modifying genotype and crypt fitness. *Cryptsim* assumes an irreversible binary model for selective mutations (each *X*_*l*_ ∈ 0 {(normal), 1 (mutated)}), and an additive model for the neutral mutations. Each selective locus *X*_*l*_, with *l* ∈ {1…*m*}, has associated selective coefficients *rl* and *sl*, which modify replication rate *b* and death rate *d*, respectively (1) and (2).

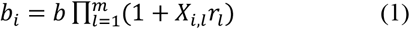

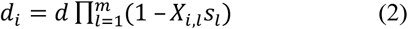

Mutation rates (μ and *v*) per locus can be relative to time or replication events, which is determined by the *a* parameter. The first option captures the effect of external mutagens (like the acidic environment in BE) while the second models errors in the DNA replication and DNA repair machinery. Equations (3) and (4) show the effect of *a* in the calculation of crypt selective (μ_*i*_) and neutral (*v*_*i*_) mutation rates per time unit for a given crypt *i*.

**Figure 1.**
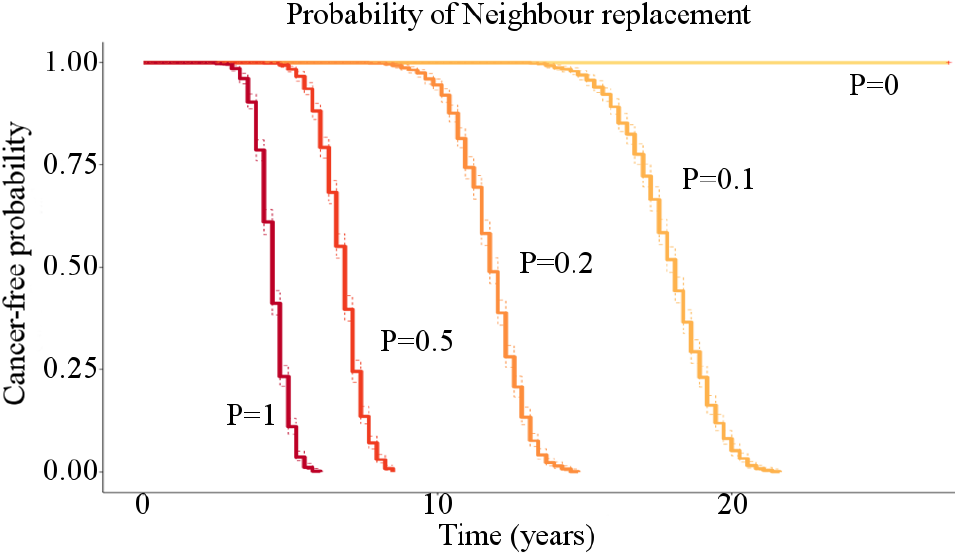
Effect on carcinogenesis of the probability of replacing a neighbor crypt (p: 1, 0.5, 0.2, 0.1, 0, indicated by different colors).

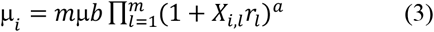

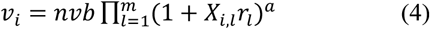

The two-dimensional spatial model represents the BE segment and consists of a *hw* (height x width) hexagonal lattice wrapped along the *h* dimension. This component allows us to consider neighbor interactions and limit the carrying capacity (i.e., maximum population size) of the simulated neoplasm; and can be either empty or full of normal crypts when the simulation starts.

Finally, a clone tree keeps track of the ancestral relationships of the different clones, their genotypes and divergence times.

### A. Strategies to overcome computational challenges

Forward simulation of a BE segment using biologically-reasonable parameters requires on the order of one billion crypt divisions, which makes this problem computationally intensive. *Cryptsim* implements a series of strategies to reduce its computational burden, among which we would like to highlight the usage of the Gillespie algorithm [11] and the calculation of genotypes on-the-fly using the clone tree. Rather than checking every crypt every time step for a rare event, we calculate the time until the next event and keep an ordered queue of events. This algorithm can be summarized in the following steps: a) sample the waiting time to the next event from an exponential distribution with rate equal to the sum of rates of all events, b) sample the event with probability proportional to its relative rate, c) choose the crypt that carries out the event at random, d) advance the sampled waiting time and carry out the event, e) update rates and iterate. The clone tree reduces memory requirements and operations by not storing genomes of each crypt (*X*_*i*_). Thus, crypts pertain to the spatial model and are linked to clones in the clone tree and rate groups in an associative array (determined by their selective genome). Mutations generate new clones in the clone tree, which also tracks their new mutation and rate group. Crypt replacements only change crypt information and their linkage with the clone tree, avoiding copying *X*_*i*_ for every replication. Clone genotypes are obtained with a simple tree traversal when needed. This clone tree is regularly pruned to remove extinct subtrees, minimizing memory requirements.

### B. Settings in which these methods are useful

As-is, this model can be applied to study evolutionary dynamics of somatic evolution of two-dimensional neoplasms organized in crypts, such as BE or the colon. However, most of the computational techniques detailed here can be applied to the study of somatic evolution in general.

### C. Availability

*Cryptsim* is implemented in Java and will be distributed under the GPL v3 license upon publication. In the meantime, contact the authors to get access to the source code.

## Acknowledgment

We thank Todd Parsons for suggesting using a Gillespie algorithm for efficient simulation.

## References

[1] F. Hvid-Jensen, L. Pedersen, A. N. Drewes, H. T. Sørensen, and P. Funch-Jensen, “Incidence of Adenocarcinoma among Patients with Barrett’s Esophagus,” N. Engl. J. Med., vol. 365, pp. 1375–1383, 2011.

[2] L. J. Esserman, I. M. Thompson, B. Reid, P. Nelson, D. F. Ransohoff, H. G. Welch, S. Hwang, D. A. Berry, K. W. Kinzler, W. C. Black, M. Bissell, H. Parnes, and S, Srivastava, “Addressing overdiagnosis and overtreatment in cancer: a prescription for change,” Lancet Oncol. vol. 15, no. 6, pp. e234–e242, 2014.

[3] A. Fortunato, A. M. Boddy, D. Mallo, A. Aktipis, C. C. Maley, and W. J. Pepper, “Natural Selection in Cancer Biology: From Molecular Snowflakes to Trait Hallmarks,” Cold Spring Harb. Perspect. Med., 2017.

[4] B. J. Reid, X. Li, P. C. Galipeau, and T. L. Vaughan, “Barrett’s oesophagus and oesophageal adenocarcinoma: time for a new synthesis,” Nat. Rev. Cancer, vol. 10, no. 2, pp. 87–101, 2010.

[5] C. C. Maley, P. C. Galipeau, J. C. Finley, V. J. Wongsurawat, X. Li, C. A. Sanchez, T. G. Paulson, P. L. Blount, R. Risques, P. S. Rabinovitch, and B. J. Reid, “Genetic clonal diversity predicts progression to esophageal adenocarcinoma,” Nat. Genet. vol 38, pp. 468–473, 2006.

[6] L. M. Merlo, N. A. Shah, X. Li, P. L. Blount, T. L. Vaughan, B. J. Reid, and C. C. Maley, “A comprehensive survey of clonal diversity measures in Barrett’s esophagus as biomarkers of progression to esophageal adenocarcinoma,” Cancer Prev. Res. (Phila), vol. 3, no. 11, pp. 1388–1397, 2010.

[7] P. C. Galipeau, L. J. Prevo, C. A. Sanchez, G. M. Longton, and B. J. Reid, “Clonal Expansion and Loss of Heterozygosity at Chromosomes 9p and 17p in Premalignant Esophageal (Barrett’s) Tissue.” J. Natl. Cancer Inst., vol. 91, no. 24, pp. 2087–2095, 1999.

[8] S. J. Leedham, S. L. Preston, S. A. McDonald, G. Elia, P. Bhandari, C. Poller, R. Harrison, M. R. Novelli, J. A. Jankowski, and N. A. Wright, “Individual crypt genetic heterogeneity and the origin of metaplastic glandular epithelium in human Barrett’s oesophagus,” Gut, vol. 57, no. 8, pp. 1041–1048, 2008.

[9] R. Kostadinov, C. C. Maley, and M. K. Kuhner, “Bulk Genotyping of Biopsies Can Create Spurious Evidence for Heterogeneity in Mutation Content,” PLOS Comp. Biol., vol. 12, no. 4, 2016.

[10] D. Hanahan, “Hallmarks of Cancer: The Next Generation,” Cell, vol. 144, no. 5, pp. 646–674, 2011.

[11] D. T. Gillespie, “A general method for numerically simulating the stochastic time evolution of coupled chemical reactions,” J. Comp. Phys., vol. 22, no. 4, pp. 403–434, 1976.

